# Dynamics of *Borrelia Burgdorferi* Invasion and Intravasation in a Tissue Engineered Dermal Microvessel Model

**DOI:** 10.1101/2022.07.10.499449

**Authors:** Zhaobin Guo, Nan Zhao, Tracy D. Chung, Anjan Singh, Ikshu Pandey, Linus Wang, Xinyue Gu, Aisha Ademola, Raleigh M. Linville, Utpal Pal, J. Stephen Dumler, Peter C. Searson

## Abstract

Lyme disease is a tick-borne disease prevalent in North America, Europe, and Asia. Dissemination of vector-borne pathogens, such as *Borrelia burgdorferi* (*Bb*), results in infection of distant tissues and is the main contributor to poor outcomes. Despite the accumulated knowledge from epidemiological, *in vitro*, and in animal studies, the understanding of dissemination remains incomplete with several important knowledge gaps, especially related to invasion and intravasation at the site of a tick bite, which cannot be readily studied in animal models or humans. To elucidate the mechanistic details of these processes we developed a tissue-engineered human dermal microvessel model. Fluorescently-labeled *Bb* (B31 strain) were injected into the extracellular matrix (ECM) of the model to mimic tick inoculation. High resolution, confocal imaging was performed to visualize *Bb* migration in the ECM and intravasation into circulation. From analysis of migration paths we found no evidence to support adhesin-mediated interactions between *Bb* and components of the ECM or basement membrane, suggesting that collagen fibers serve as inert obstacles to migration. Transendothelial migration occurred at cell-cell junctions and was relatively fast, consistent with *Bb* swimming in ECM. In addition, we found that *Bb* alone can induce endothelium activation, resulting in increased immune cell adhesion but no changes in global or local permeability. Together these results provide new insight into the minimum requirements for dissemination of *Bb* at the site of a tick bite, and highlight how tissue-engineered models are complementary to animal models in visualizing dynamic processes associated with vector-borne pathogens.

**Significance Statement:** Using a tissue-engineered human dermal microvessel model we reveal new insight into the invasion and intravasation of *Borrelia burgdorferi* (*Bb*), a causative agent of Lyme disease in North America, following a tick bite. These results show how tissue-engineered models enable imaging of dynamic processes that are challenging in animal models or human subjects.

## Introduction

Lyme disease is prevalent in North America, Europe, and Asia (1), and is the most common vector-borne disease in the US (2). The most recent data from the CDC estimates 476,000 new cases every year (3, 4). While antibiotic treatment is effective, some individuals experience symptoms for months or years following treatment (5). Lyme disease can lead to health problems associated with the skin, joints, central nervous system (neuroborreliosis) and, to a lesser extent, the heart (5, 6).

Lyme disease in North America is caused primarily by the spirochete *Borrelia burgdorferi* (*Bb*) and is transmitted to humans by a bite from an infected tick (1). Dissemination of vector borne pathogens, such as *Bb*, involves several critical steps, including inoculation in the dermis, proliferation and migration in the local tissue, intravasation into circulation or the lymphatic system, and extravasation into and colonization of distant tissues and organs (1, 6). Most of our knowledge about dissemination comes from analysis of tissue samples in mouse models, e.g. enumeration of *Bb* at the inoculation site or in other tissues. Since the processes associated with dissemination are dynamic, visualization is key to elucidating mechanisms. A relatively small number of intravital microscopy (IVM) studies in mouse models have been key in beginning to unravel the details of *Bb* extravasation (7–17). These studies have been complemented by novel studies of spirochete adhesion in flow chambers (18), 3D migration studies (19), and membrane feeding assays (20). While these studies have been able to establish important links to results from microbiological studies, the biological and microenvironmental factors that regulate *Bb* dissemination, especially related to intravasation, remain poorly understood.

To address the knowledge gaps associated with *Bb* invasion and intravasation during the early sub-acute phase of infection, we developed a tissue-engineered human dermal microvessel model in a type I collagen extracellular matrix (ECM), the main structural component of the loose connective tissue of the dermis and matched to its stiffness (21–23). GFP-labelled *Bb* (B31 strain) were inoculated in the ECM to mimic a tick bite. High resolution, confocal imaging was then performed to visualize *Bb* migration in the ECM and intravasation into circulation. Based on *in vitro* studies it has been postulated that *Bb* migration is directed by chemoattractants, and that both invasion and interactions with the endothelium are regulated by adhesins (19, 24–26). We show that in a 3D microvessel model, there is no evidence for chemoattraction, and that adhesins do not play a significant role in migration. *Bb* in the 3D matrix exhibited the same modes of motion as reported in 2D: forwards, backwards, and stationary (27). However, we show that the distribution of reversal events is significantly increased in 3D, which we attribute to the collagen fibers that serve as obstacles. We show that intravasation occurs at cell-cell junctions following a trial and error searching process. While *Bb* swimming through cell-cell junctions can occur with little resistance, in many cases, the rear end of the spirochete cell body is transiently tethered to the endothelium (for up to 100 s) prior to release into circulation. Finally, we show that *Bb* can induce endothelium activation in the absence of systemic inflammatory cytokines while maintaining normal barrier function. While these results remain to be further refined by independent measurements (e.g. knockout studies of *Bb* adhesins) and in more complex models (e.g. inclusion of tick saliva), we show how the ability to independently control variables in a reductive tissue-engineered human model can advance our understanding of the complexity of *Bb* dissemination in humans.

## Results

### Creation of B. burgdorferi local invasion model

The critical steps in tick-borne pathogen dissemination, e.g. invasion, intravasation, arrest, and extravasation, take place at vascular interfaces (28). Therefore, we developed a tissue engineered dermal microvessel model to enable visualization of invasion and intravasation of *B. burgdorferi* (*Bb*) in a platform that enables independent control of key experimental variables (**Fig. 1a – d**). The dermal microvessel was formed by seeding human dermal microvascular endothelial cells (HDMECs) into a 150 μm diameter channel in a type I collagen matrix (**Fig. 1a, b**). Following adhesion and spreading, the HDMECs form a confluent monolayer within 24 hours. To simulate tick inoculation, fluorescently-labeled *Bb* (B31-A3 GFP strain) were injected into a small cavity ~900 μm away from the microvessel (**Fig. 1c, d** and **Fig. S1a**) at 48 h after seeding (**Fig. S1b**). During experiments, the dermal microvessel was perfused with endothelial cell medium at a shear stress of ~4 dyne cm^-2^, corresponding to typical values in post-capillary venules.

**Figure 1.**
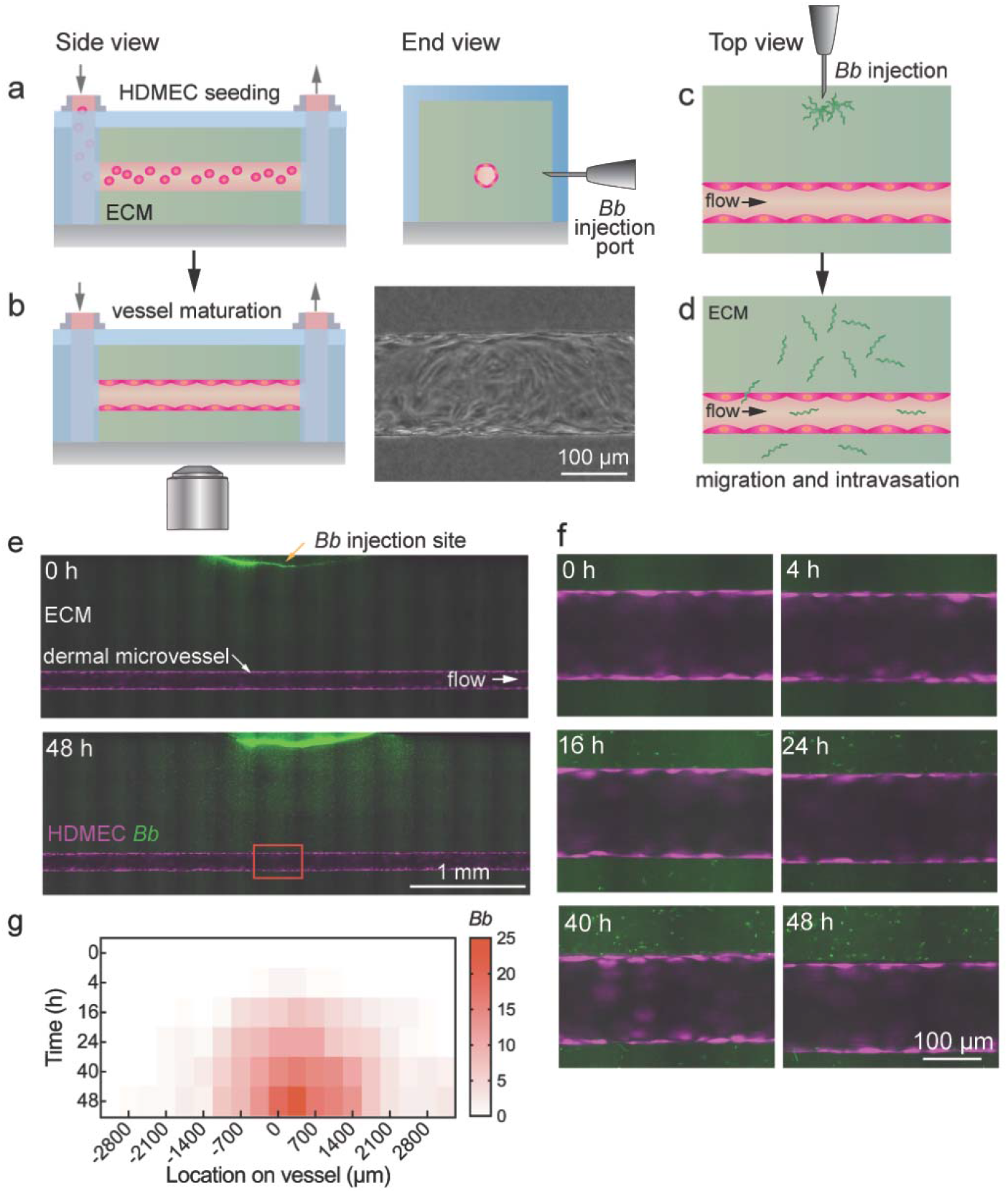
Invasion of *B. burgdorferi (Bb)* in a tissue-engineered dermal microvessel model. **(a)** Human dermal microvascular endothelial cells (HDMECs) seeded into the microfluidic device in collagen type I extracellular matrix. **(b)** HDMECs form a confluent monolayer after 2 days under shear stress (4 dyne cm^-2^). **(c)** Inoculation of GFP-tagged *Bb* into the dermal microvessel. **(d)** Following inoculation, *Bb* migrate from the injection site. **(e)** Fluorescence images of dermal microvessels at 0 h and 48 h following inoculation with fluorescently-labeled *Bb* (green). HDMECs (magenta). Flow was maintained at 4 dyne cm^-2^. **(f)** Fluorescence images in the vicinity of the inoculation site (see red box in panel (e)) over time. **(g)** Heat map showing quantification of *Bb* in the perivascular region along the vessel over time. Imaging was performed at the equatorial plane of the microvessel with a depth of field of ~4.3 μm. The *Bb* concentration was obtained in a region within 20 μm of the endothelium at the equatorial plane on both sides of the vessel (i.e. anterior and posterior to the inoculation site). The location along the microvessel is relative to the point closest to the inoculation site and extends about 3 mm in both upstream and downstream directions.

Following inoculation (**Fig. 1c)**, *Bb* migrate within the ECM (**Fig. 1d**) and were observed beyond the microvessel within 1 h. At 48 h following inoculation, a large number of *Bb* were observed localized around the microvessel (**Fig. 1e, f**). To assess differences in the vicinity of the microvessel, we define the region within 20 μm of the microvessel, approximating the length of a typical *Bb* spirochete (27), as the perivascular region, and the region > 20 μm from the endothelium as bulk ECM. The concentration of *Bb* in the perivascular region at the equatorial plane (**Fig. S1c**) increased over time, and at all time points was highest at the point closest to the inoculation site and decreased laterally in the upstream and downstream regions (**Fig. 1g**). In addition, the density of *Bb* on the anterior side of the vessel was higher than that on the posterior side relative to the inoculation site (**Fig. S1d, e and Supplementary Note S1**).

### Characterization of B. burgdorferi migration

The migration paths of individual *Bb* in the ECM were tracked by determining the location of the midpoint along the *Bb* spirochete (**Fig. 2a and Video S1**). Each migration path consisted of a series of individual segment vectors describing distance and angle at each time point (Δt = 1 s). Tracking was limited to *Bb* with a residence time within the focal plane of at least 20 s. The median tracking time was ~100 s and, in total, we recorded 10,131 segments for 43 individual *Bb*. From analysis of the migration paths (**Fig. 2b**), we found negligible net displacement perpendicular (along y-axis, −0.60 ± 23.5 μm) or parallel (along x-axis, 1.11 ± 23.3 μm) to the microvessel (**Fig. 2c**), showing that migration was random and that factors such as chemotaxis or interstitial flow did not play a significant role in *Bb* migration.

**Figure 2.**
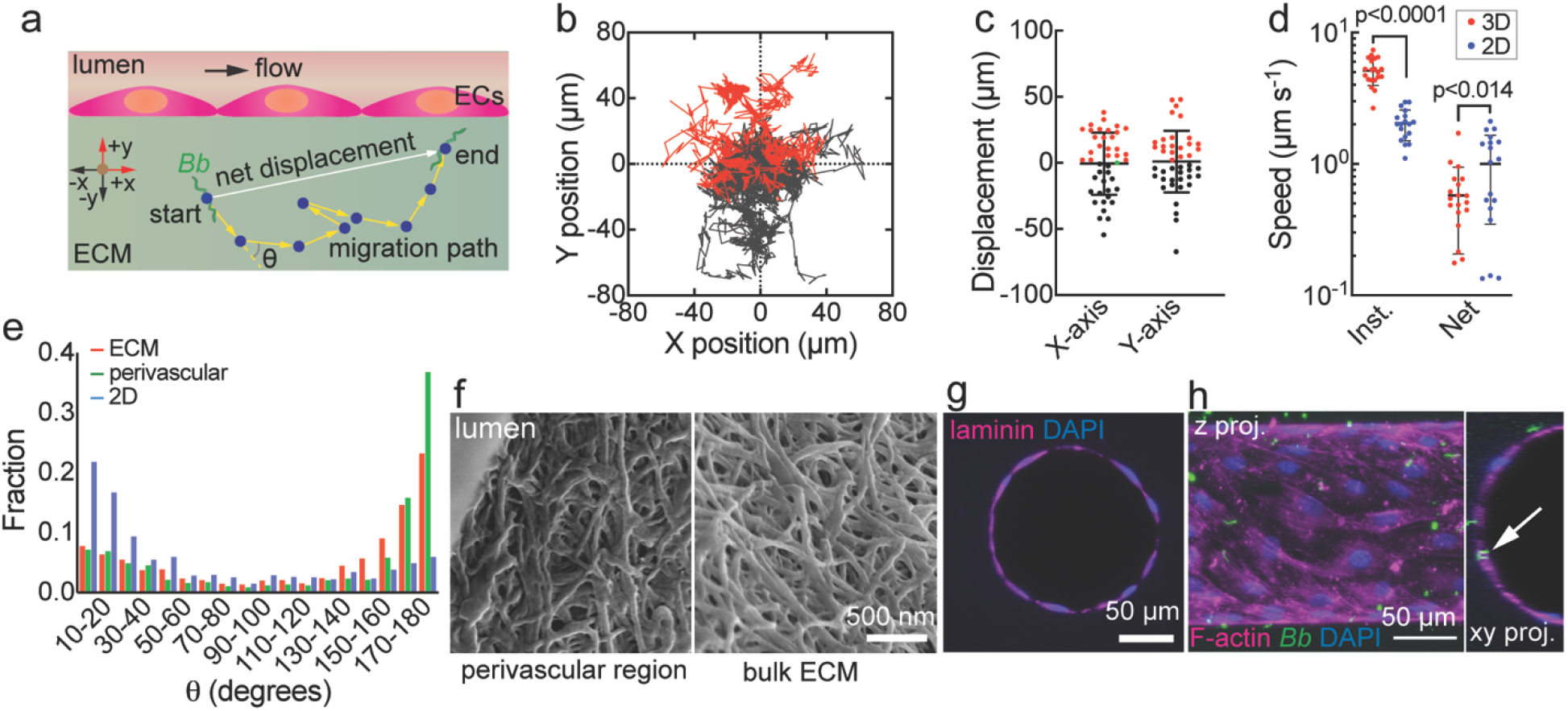
*Borrelia burgdorferi (Bb)* migration in ECM and at the vessel-ECM interface. **(a)** Schematic illustration showing the migration path of a single *Bb* in the matrix. Tracking was limited to cases where *Bb* were resident within the focal plane for at least 20 s. The average tracking time for a single *Bb* was 117 ± 138 s. The location of a *Bb* in each image was determined from the midpoint along the length of the spirochete cell body (denoted by blue circles). The yellow arrows represent the segment vectors between each image. The instantaneous speed was determined from the difference in position between successive images (Δt = 1s). **(b)** Overlay of migration paths for *Bb* in the ECM within the focal plane (n = 43). The start points for all migration paths are set to the origin. Red and black lines indicate positive or negative migration direction in the y-axis: +y is towards the microvessel, and +x is in the downstream direction with respect to flow. **(c)** The net displacement of *Bb* perpendicular (y-axis) and parallel (x-axis) to the microvessel (n = 43). Positive (red) and negative (black) displacement along x- or y-axis (see panel (a)), green indicates zero net displacement. **(d)** Net and instantaneous speed for *Bb* migration in the ECM (n = 20) and in the 1% methylcellulose solution (n = 18). Note that a data point for 3D net speed (4.75×10^-3^ μm s^-1^) is not shown but was included in statistical analysis. **(e)** Distribution of the angles (θ) between two consecutive vectors along the migration path in the ECM (n = 2,081 segments for 20 *Bb*), the perivascular region (n = 1,948 segments for 5 *Bb*), and in 1% methylcellulose solution (n = 3,470 segments for 18 *Bb*). **(f)** SEM images of type I collagen ECM from the perivascular region and bulk ECM. **(g)** Immunofluoresence staining shows a well-defined layer of the basement membrane protein laminin-α4 at the vessel-ECM interface. **(h)** Confocal images of a dermal microvessel 24 h after inoculation showing transmigration of a single *Bb* (white arrow).

The instantaneous speed for *Bb* migration was determined from the elapsed time for each segment along the migration path. For comparison, we performed experiments with *Bb* in 1% methylcellulose solution (denoted as 2D) (**Video S2**), which is commonly used to study *Bb* “swimming” *in vitro* (27, 29–32). The instantaneous speed in 3D was significantly higher than in 2D (5.14 ± 1.18 μm s^-1^ vs. 2.07 ± 0.51 μm s^-1^, p < 0.0001) (**Fig. 2d**). In contrast, the net speed (the net displacement divided by the elapsed time for each migration path) in 3D was significantly lower than that in 2D (0.58 ± 0.37 μm s^-1^ vs. 1.01 ± 0.65 μm s^-1^, p < 0.05). The instantaneous speed in perivascular region was 3.39 ± 7.91 μm s^-1^, which is slightly lower than in the ECM (5.16 ± 3.75 μm s^-1^, p < 0.001) (**Fig. S2a**)

Next, to assess the mode of *Bb* migration, we determined the angle (θ) between consecutive segments along the migration paths (**Fig. 2a**). In all 3 conditions (ECM, perivascular region, and 2D), the distribution of θ was asymmetric: in 2D the distribution was biased towards small θ (relatively straight trajectories), whereas in the perivascular region or in the ECM, the distributions were biased towards large θ (large changes in direction) (**Fig. 2e**). In general, θ is mainly distributed between 150°−180° in 3D, compared to 0 - 30° in 2D (**Fig. 2e**). In 3D, backwards motion (θ > 90°) accounts for about 2/3 of segments (65.4% in ECM, 68.8% in the perivascular region), whereas in 2D forwards motion (θ ≤ 90°) accounts for about 2/3 of segments (forward 69.2%) (**Fig. S2b**). Further analysis confirmed that *Bb* in 3D exhibited less persistent forwards motion (**Fig. S2d, e and S2h**) but more back-and-forth motion (**Fig. S2f, g and S2h**) compared to 2D. The frequency of persistent back-and-forth motion was slightly higher in the perivascular region compared to the ECM (see **Supplementary Note 2**). The larger frequency of back-and-forth motion explains why the net velocity is lower in 3D compared to 2D. In 3D, the collagen fibers provide obstacles to forward motion, resulting in a larger number of direction changes and more persistent back-and-forth motion. The higher instantaneous velocity in 3D is due to the lower viscosity of the interstitial fluid compared to 2D. All of the “stationary” segments (vector length ≤ 0.4 μm) were excluded from analysis, but only account for a small fraction of time in both 2D and 3D. Stationary states represented 1.1 % of events (23 of 2,104) in the ECM, 4.3 % (87 of 2,035) in the perivascular region, and 3.0 % (106 of 3,576) in 2D. In addition, most stationary events were transient (< 2 s) (**Fig. S2c**): longer stationary events represented ~1% of all stationary events and less than 0.1% of all segments. In the perivascular region the instantaneous speed parallel to the endothelium was lower (2.62 ± 1.86 μm s^-1^) compared to perpendicular to the endothelium (4.37 ± 2.89 μm s^-1^) (**Fig. S3a**), and the fraction of stationary events was higher and more persistent (**Fig. S3b, c and Supplementary Note 3)**. This could be associated with an increase in viscosity associated with basement membrane proteins around the endothelium.

We next characterized the microstructure of the collagen matrix in the bulk and the perivascular region. Scanning electron microscope images of lyophilized gels (**Fig. 2f**) were very similar to decellularized vascular grafts (33) with fiber diameters of 100 - 200 nm. The porosity of hydrated gels with the same collagen concentration is ~ 95 % (34, 35), and the pore size is estimated to be several hundred nanometers. There was no visible difference in ECM structure in the perivascular region compared to the bulk ECM (**Fig. 2f**), suggesting that the difference in migration speeds between ECM and the perivascular region is not related to ECM density. Immunofluorescence images of laminin confirmed the presence of a layer of basement membrane around microvessels (**Fig. 2g**). When *Bb* contacted the microvessel endothelium, some interactions resulted in insertion into the endothelium (**Fig. 2h**), a prerequisite for intravasation.

### Transendothelial migration and intravasation

Although the number of *Bb* in the vascular region following inoculation was relatively high (**Fig. 1g,**), intravasation events were rare. Intravasation events were analyzed if the *Bb* remained in the field of view at the equatorial plane of the microvessel during the initial interaction with the endothelium, transmigration, and intravasation. Across 19 analyzable intravasation events, two general mechanisms were observed: direct and indirect intravasation. In the direct mechanism, *Bb* contacted the microvessel and immediately transmigrated (**Fig. 3a,b**), a process occurring over a few seconds. Microvessels with fluorescently labeled HDMECs showed that *Bb* inserted directly into the cell-cell junction prior to intravasation (**Fig. 3a** and **Movie S3**). In the indirect mechanism, *Bb* arrived at the endothelium, but underwent one or more cycles of back-and-forth motion, either parallel or perpendicular to the microvessel/ECM interface, prior to transmigration (**Fig. 3c**, **Fig. S4a-d**; **Movies S4 - S6**). The lateral displacement during this process was typically less than the length of an endothelial cell body. From analysis of videos with fluorescently-labeled endothelial cells (**Fig. 3a,c** and **Fig. S4e**), we confirmed that all transmigration events occurred at cell-cell junctions. In summary, over 19 events, 5 displayed direct transmigration, while 14 showed at least 1 cycle of back-and-forth motion while locating a cell-cell junction. Once most of the spirochete cell body had crossed the endothelium, *Bb* often remained transiently tethered for up to 100 s to the endothelium prior to complete intravasation (**Fig. 3d,e**).

**Figure 3.**
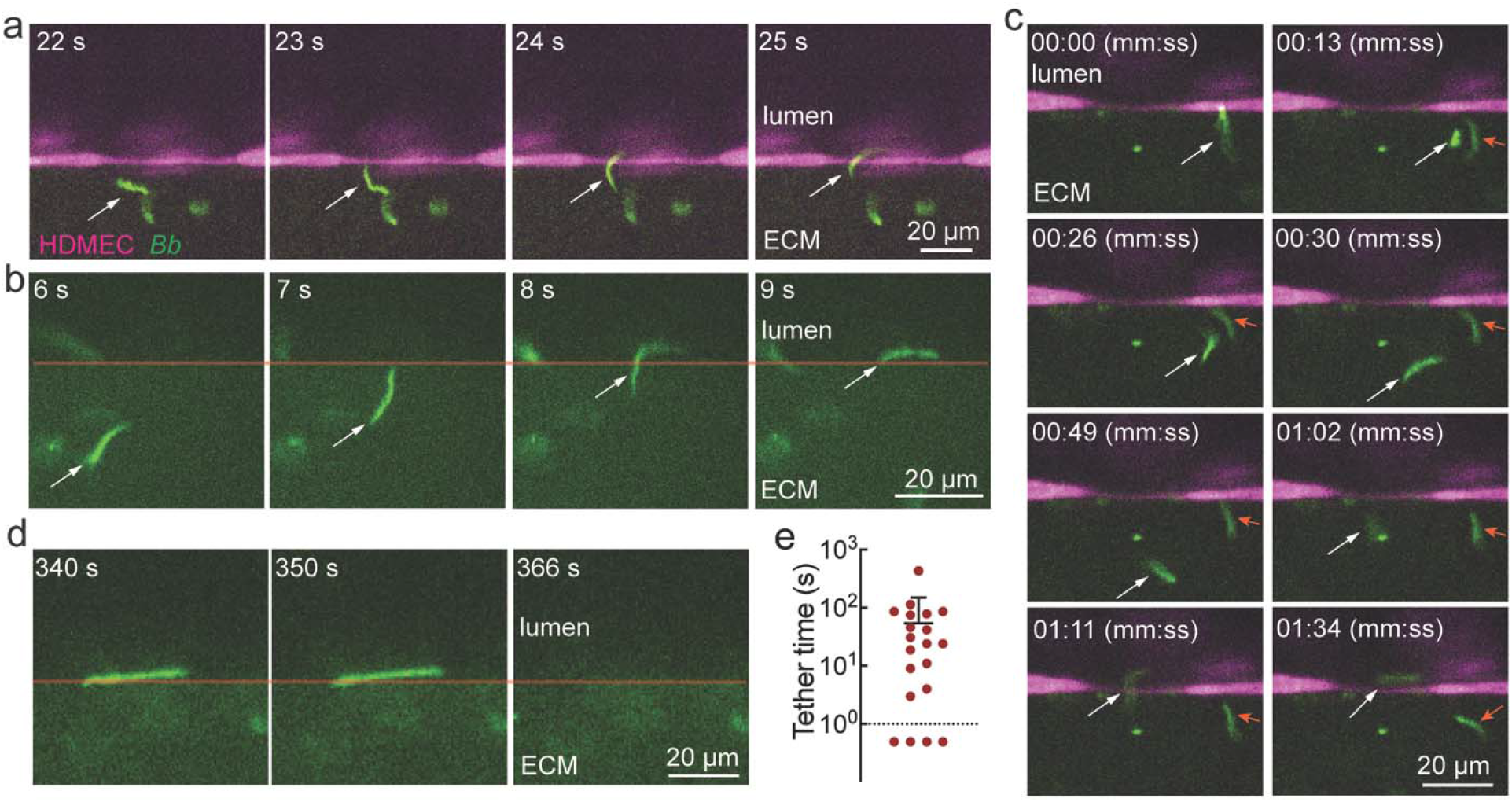
Dynamics of *Borrelia burgdorferi (Bb)* transmigration. **(a)** Example of direct intravasation where the initial contact of a *Bb* (white arrow) with the endothelium is at a cell-cell junction. Transmigration occurs over 2 s. HDMECs (magenta), *Bb* (green). **(b)** Example of direct intravasation. The red line indicates the position of the endothelium (obtained from phase images, not shown). **(c)** Example of indirect intravasation where a *Bb* (white arrow) arrives at the microvessel, touches the endothelial cell body several times prior to migration to a nearby cell-cell junction and intravasation. A second *Bb* (red arrow) also shows back and forth motion on arriving at the endothelium. **(d)** Transient tethering during transmigration. Although most of the *Bb* cell body has transmigrated into the lumen, the rear remains transiently tethered to the endothelium for several seconds prior to intravasation. **(e)** Quantification of tethering time following transmigration. For 4 events there was no transient tethering.

From imaging at 48 h post-inoculation, the rate (N·h^-1^) of *Bb* contacting the endothelium was ~ 10^9^ per hour, of which 2.33 ± 0.70% (n = 3 microvessels) resulted in intravasation. Since the region of imaging corresponds to ~1% of the entire microvessel, we estimate an intravasation rate of around 250 per hour from the simulated tick bite. Assuming a *Bb* lifetime in circulation of ≤ 1 h, this corresponds to a blood concentration (assuming 5 L) of 0.05 per mL. This is in agreement with the fact that *Bb* are rarely detected in blood samples of individuals with Lyme disease where the upper limit would be < 1 in a 10 mL blood sample or 0.1 per mL (36).

### Bb and endothelium activation

To assess the effects of *Bb* on the endothelium, we perfused microvessels with monocytic (THP-1 cells) or promyelocytic (HL-60) cells 48 h following inoculation, and quantified the density of adherent cells following wash-out of suspended cells (**Fig. 4a**). Microvessels inoculated with *Bb* showed a significant increase in the number of adherent leukocytes (**Fig. 4a, b**) suggesting that *Bb* alone, in the absence of extrinsic inflammatory molecules, can activate endothelial cells and elicit an inflammatory immune response. Immunofluorescence images confirmed upregulation of the adhesion molecule ICAM-1 by 3.68 ± 2.17-fold (**Fig. 4c-e**) following inoculation with *Bb* compared to its corresponding vehicle.

**Figure 4.**
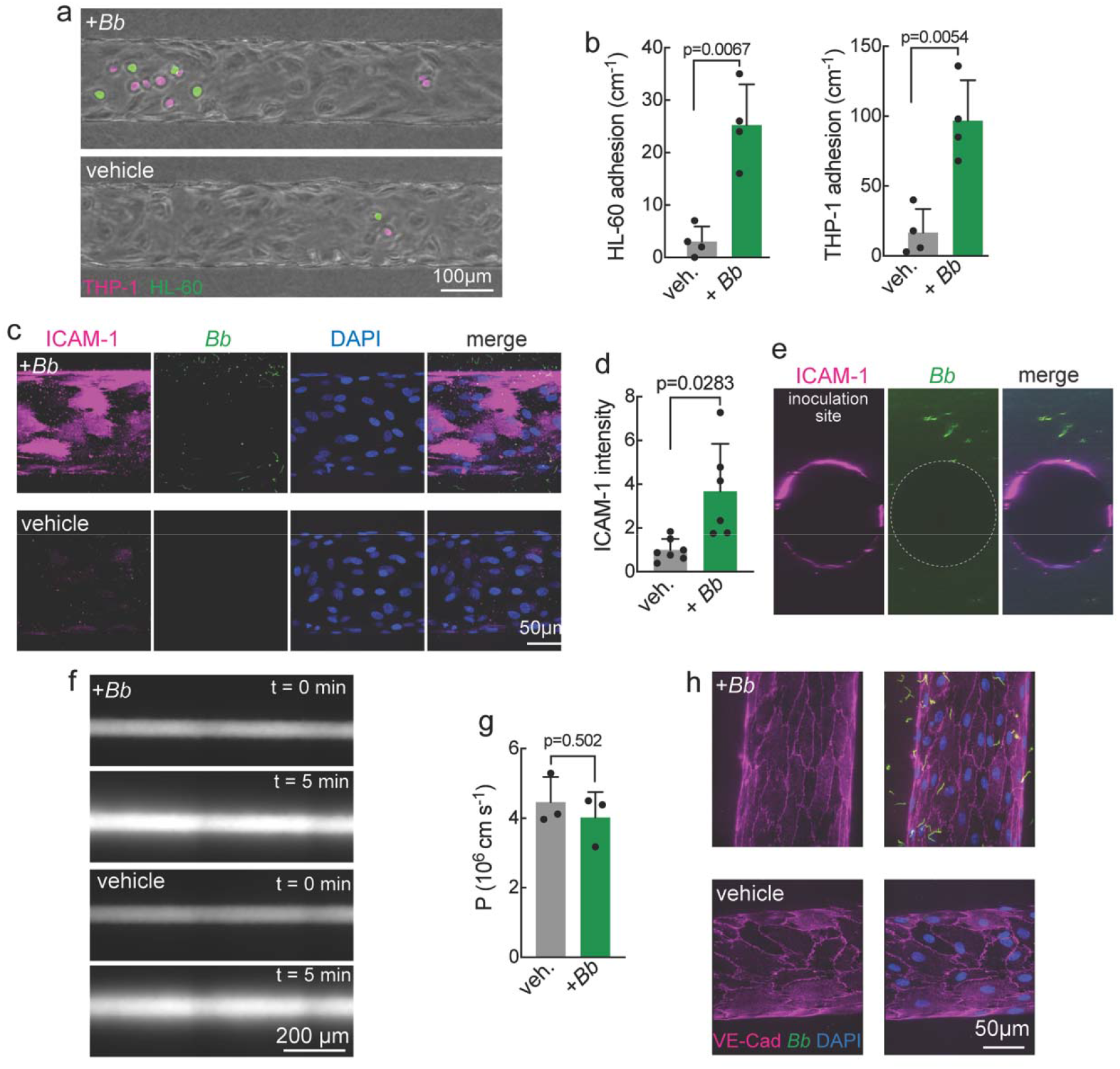
Inoculation with *Bb* induces endothelium activation but does not influence global or local barrier function. **(a)** Fluorescence images of HDMEC microvessels perfused with leukocytes (1 × 10^6^ mL^-1^ THP-1 and 1 × 10^6^ mL^-1^ HL-60) for 10 min at 48 h following inoculation with *Bb* or vehicle. Imaging was performed 15 min following perfusion with leukocytes. THP-1(red), HL-60 (green). **(b)** The number of adherent THP-1 (left panel) and HL-60 (right panel) cells was significantly higher in the presence of *Bb* (n = 4 independent microvessels). **(c)** Representative confocal z-axis maximum intensity projection images of ICAM-1 (magenta) and DAPI (blue) in microvessels 48 h following inoculation with *Bb* (green) or vehicle. **(d)** ICAM-1 fluorescence intensity measured from confocal z-axis maximum intensity projection images of microvessels 48 h following inoculation with *Bb* (green) or vehicle. *Bb* inoculation: n = 6 microvessels. Vehicle: n = 7 microvessels. Each point represents an independent experiment relative to the averaged value from vehicle controls. **(e)** Maximum intensity projection image (top panel) and slice at the equatorial plane (bottom panel) of ICAM-1(deep red) inoculation with *Bb* (green). The inoculation site was towards the top of the microvessels in this orientation. **(f)** Fluorescence images of dermal microvessels 48 h after inoculation with *Bb* or vehicle during perfusion with 2 MDa dextran. **(g)** Permeability of 2 MDa dextran was the same 48 h following inoculation with *Bb* or vehicle. **(h)** Immunofluorescence images of VE-cadherin at 48 hours following inoculation with *Bb* or vehicle.

To determine whether *Bb* induced changes in barrier function, we measured microvessel permeability by perfusing with fluorescently-labeled 2 MDa dextran (**Fig. 4f**). The permeability was ~ 4 ×10^-6^ cm s^-1^, and there was no difference between conditions (**Fig. 4g**). Since the size of 2 MDa dextran is around 50 nm (37), these results suggest that the paracellular gaps are sufficiently large to allow *Bb* migration. Immunofluorescence imaging of VE-cadherin in microvessels showed well-formed adherens junctions, with no difference between *Bb* and vehicle (**Fig. 4h**).

## Discussion

Dissemination of vector-borne pathogens involves several critical steps, including inoculation in the dermis, proliferation and migration in the local tissue, intravasation into circulation or the lymphatic system, and extravasation and colonization of distant tissues and organs (1, 6). Much of our current knowledge of the interactions of pathogens with the vascular system comes from IVM studies in mouse models (7–17) and *in vitro* models (e.g. Boyden chamber). 2D Transwell models capture some aspects of the dynamics of dissemination, but are reductive and do not allow real-time imaging. 3D tissue-engineered models vary in complexity and physiological relevance, but provide a diverse toolkit for the study of vascular phenomena (38, 39). We have developed a 3D tissue-engineered dermal microvessel model to visualize invasion and intravasation of *Borrelia burgdorferi* (*Bb*), the causative agent of Lyme disease (40), following inoculation into the ECM. Tissue-engineered models recapitulate the cylindrical vessel geometry, physiological flow rates, and incorporate human endothelial cells in contact with basement membrane embedded within an ECM.

### Bb migration in ECM

Following inoculation in the human dermal microvessel model, *Bb* exhibited the three modes of motion observed *in vitro*: forwards motion, backwards motion, and a stationary state (27). The instantaneous speed in ECM (5.1 μm s^-1^, **Fig. 2d**) is very close to values reported *in vitro* in gelatin hydrogels (3.5 - 5 μm s^-1^) (41), and *in vivo* in the ear dermis in a mouse model (up to 4.0 μm s^-1^) (32, 41, 42), but faster than *Bb* swimming in 2D (2.1 μm s^-1^, **Fig. 2d**) (viscosity of 1% methylcellulose: ~ 200 mPa·s). The distribution of the modes of motion were different between conditions. In 2D, forward motion was persistent and there was a relatively small fraction of reversal events. In contrast, in 3D there was a much larger fraction of reversal events, likely due to the collagen fibers which act as obstacles for forward motion. The average porosity of 3 - 7 mg mL^-1^ type I collagen gels is around 95% (34, 35). For an average fiber diameter of ~150 nm (**Fig. 2f**), this corresponds to an average spacing of around 400 nm. Therefore, during 1 s (imaging frequency), a *Bb* is expected to encounter more than 10 collagen fibers, some of which may result in large displacement angles between segments.

*Bb* express adhesins on their outer membranes, including proteins such as BB0406 (43–45) and BmpA (46), which bind to the basement membrane protein laminin, and BBK32 which binds to the ECM protein fibronectin (47, 48). While *BB0406*^-^ and *BBK32*^-^ mutant *Bb* result in reduced numbers of spirochetes in tissues in mice (43), their exact role in invasion and intravasation remains unknown. Our results suggest that adhesins do not play a significant role in *Bb* migration in the ECM. The instantaneous velocity in the collagen I matrix is faster than in methylcellulose suggesting that any binding interactions with fibronectin in the ECM is negligible. Studies of BBK32 binding to fibronectin found *K_D_* = 0.019 μM L^-1^ and *k*_off_ = 0.10 ± 0.054 s^-1^, corresponding to an average residence time of around 10 s (49), much longer than the average duration of stationary states found here (≤ 1s). Although we observed a decrease in instantaneous velocity in the perivascular region, this difference is only about two-fold and is not consistent with the hypothesis that immobilization of *Bb* by laminin binding in the basement membrane is a key step in intravasation. Studies of BB0406 binding with laminin found *K*_D_ = 0.4 μM L^-1^ and *k*_off_ = 0.31 ± 0.021 s^-1^ (43), an average residence time of around 3 s. Under static conditions, the residence times would be expected to result in a population of relatively long-lived stationary states during migration in extracellular matrix or basement membrane. Since the duration of stationary states is relatively short, it is likely that the momentum generated by the flagella motors is sufficiently large to overcome the binding to ECM components. The large fraction of reversal events in the perivascular region (~40% of segments > 170°) is likely due to the presence of the endothelial cell bodies, which provide large obstacles to migration.

### Transmigration and intravasation

We observed that intravasation occurs at cell-cell junctions via direct or indirect transendothelial migration. If the initial point of contact with the endothelium was at a cell-cell junction, then the *Bb* continued to migrate through the junction. If the first trial failed, the *Bb* exhibited back-and-forth motion and either located a cell-cell junction, or completely changed direction to a nearby location until a cell-cell junction was found. These results are consistent with the hypothesis that *Bb* and other spirochetes (e.g. *Leptospira)* use back-and-forth motion as a trial and error method to circumvent obstacles in tissues (32, 50). We found no directional bias in the migration of *Bb* (**Fig. 2b,c**), implying that the initial point of contact at the endothelium was purely random. However, this remains to be verified since we could not determine the fraction of contact events that resulted in intravasation due to *Bb* migration out of the field of view.

Although transmigration at cell-cell junctions was relatively fast, *Bb* often remained tethered to the endothelium at their rear for up to 100 s prior to intravasation. A possible explanation is that insertion into the lumen triggers reversal of the rear flagella motors to provide a resistance to the fluid shear force in the lumen. *Bb* are typically around 0.3 μm in diameter and 10 - 20 μm in length (51). In the examples shown here, transmigration occurred over a few seconds suggesting that the process involves swimming through the cell-cell junctions in the endothelium in the absence of biochemical interactions. At a swimming speed to 5 μm s^-1^, a 10 - 20 μm long *Bb* would take 2 - 4 s to cross the endothelium, consistent with the transmigration time observed here.

In summary, live cell imaging (**Fig. 3** and **Movie S4-S7**), fluorescence profiles (**Fig. S4e**), and the fast transit time provide strong evidence in support of transmigration occurring at cell-cell junctions.

### Influence of Bb on the endothelium

Local inflammation at the site of a tick bite is a hallmark of early Lyme disease (erythema migrans) (52), and is characterized by upregulation of cytokines and chemokines associated with recruitment and activation of immune cells (53). Here we found that *Bb* alone can induce activation of the endothelium resulting in increased expression of adhesion molecules (e.g. ICAM-1) and leukocyte adhesion. These results show that *Bb* can elicit an inflammatory response in the endothelium even in absence of resident immune cells, stromal cells, or tick saliva.

Despite activation of the endothelium following inoculation with *Bb*, we found no change in the global permeability to 2 MDa dextran (hydrodynamic size 50 nm), and no evidence of focal leaks associated with local disruption of cell-cell junctions. These results suggest that barrier dysfunction is not required for intravasation and that *Bb* can transmigrate by swimming through normal gaps in cell-cell junctions (38, 54).

In summary, we developed a tissue-engineered dermal microvessel platform to study processes associated with invasion and intravasation of *Borrelia burgdorferi* (*Bb*), the causative agent of Lyme disease. High resolution, confocal imaging was employed to visualize *Bb* migration in the ECM and invasion of the dermal microvessel model. Using this model we report several key findings. (1) Migration was random with no directional bias implying that there was no chemoattraction to blood vessels. (2) We confirmed the same modes of migration as observed in 2D swimming: forwards, backwards, and stationary. (3) The distribution of angles between segments revealed a much higher fraction of reversal events in 3D, likely due to encountering collagen fibers during migration. (4) There was no evidence to support adhesin-mediated interactions between *Bb* and components of the ECM or basement membrane, suggesting that collagen fibers serve as inert obstacles to migration. (5) Transendothelial migration occurred at cell-cell junctions: initial contact with the endothelium away from cell-cell junctions resulted in cycles of back-and-forth motion or migration to a cell-cell junction. (6) Intravasation occurred over several seconds, consistent with *Bb* swimming through the cell-cell junctions. (7) Following transmigration, transient tethering of the rear end of the spirochete cell body occurred in some cases, with residence times up to 100 s. (8) *Bb* alone can induce endothelium activation, resulting in increased immune cell adhesion but no changes in global or local permeability. Together these results advance our understanding *Bb* dissemination at a tick bite, and highlight how tissue-engineered models are complementary to animal models in enabling a reductive approach to addressing key mechanistic question in a model with human cells. The reductive approach has many advantages in studying complex dynamic processes, however, our results remain to be refined (e.g. with adhesin knockout studies) and extended to include other important variables (e.g. tick saliva).

## Materials and Methods

### Fabrication of dermal microvessels and Borrelia inoculation

Microvessels were formed using previously published protocols (21, 38). Briefly, ~1 × 10^5^ human dermal microvascular endothelial cells (HDMECs, Lonza, CC-2543) were seeded into ~ 150 μm diameter channels in a 7 mg mL^-1^ type I collagen matrix (Corning, 354249) and perfused in endothelial cell medium (EGM-2) medium for 2 days to achieve a confluent monolayer (see *Supplementary Information* for details). Next, 1 × 10^7^ *Bb*-GFP (B31-A3 GFP strain, passage 4) were injected into a side port to simulate a tick bite, and imaging or functional testing performed in a live cell chamber at 37 °C and 5% CO_2_.

### Tracking Borrelia burgdorferi dissemination, migration, and intravasation

To track the dissemination of *Bb* from side port to the microvessel confocal imaging at single time point were acquired with 20 × magnification at 0 h, 4 h, 24 h, 40 h, 48 h after inoculation. The entire microvesssel (7,102 μm × 1,782 μm, 20 × 5 images) were scanned and the number of *Bb* in the vicinity of microvessel (□ 20 μm) were counted manually. To track the migration and intravasation events, time lapsed images were acquired at 20× magnification on a swept field confocal microscope for 1h with image rate of 1s/frame, and performed at 24 h or 48 h after *Bb* inoculation, focusing the region with intermediate *Bb* density. Endothelial cells were labelled with CellTracker Deep Red (Thermo Fisher Scientific, C34565) to visualize the endothelium during *Bb* intravasation. Tracking the migration of *Bb*, calculation of *Bb* migration speeds and angles (θ) between two consecutive vectors along the migration path that determining forward or backward moving was described in detail in the supplemental methods section.

### Functional assays of microvessel

To assess immune cell adhesion, microvessels were perfused with 1 × 10^6^ HL-60 cells (ATCC) and 1 × 10^6^ THP-1 cells (ATCC) for 10 minutes at a shear stress of 0.2 dyne cm^-2^ 48 h after inoculation with *Bb* or vehicle (BSK-II medium). After washing out the non-adherent immune cells, adherent HL-60 and THP-1s were manually counted separately in each device, and the number of adherent cells normalized to the area of the microvessel.

To assess endothelium barrier function we perfused microvessels with 2 μM Alexa Fluor647-conjugated 2 MDa dextran (Thermo Fisher Scientific, cat. no. D22914) in EGM-2 medium 48 h following inoculation with *Bb* or vehicle (BSK-II medium) inoculation. Phase contrast and fluorescence images (8,107 μm × 664 μm) were acquired every 30 s for 2 minutes before and 5 minutes following perfusion with the fluorescent solutes. Permeability of microvessels was calculated from P = (r/2)(1/ΔI)(dI/dt), where r is the vessel radius, ΔI is the increase in fluorescence intensity upon initiation of perfusion of the solute, and dI/dt is the rate of increase of fluorescence increase as the solute permeates into the collagen gel. Sections from −700 to +2100 μm (relative to the inoculation site) were selected for calculation of permeability as *Bb* mainly accumulated within these regions with or without *Bb* inoculation.

### Immunocytochemistry and image analysis

48 h after *Bb* inoculation, immunocytochemistry was performed with protocol described in detail in the supplemental methods. Confocal z-stacks (0.4 μm in thickness) were acquired at 40× magnification on a swept field confocal microscope system (Prairie Technologies) and reconstruction of microvessels were assembled from approximately 400 slices. To quantify the expression level of ICAM-1 (Fig. 4g), maximal intensity projection of z stack of 400 slices was performed and fluoresce intensity was measured in Image J with background subtraction. Reported data was normalized to the averaged fluorescence intensity of microvessels (n=7) from control group.

### Statistics

All experimental values are reported as mean ± standard deviation (S.D.). A student’s unpaired t-test (two-tailed with unequal variance) was used for comparison of two groups. Differences were considered statistically significant for p < 0.05.

## Author Contributions

P.S., Z.G., U.P., J.D. conceived the original idea. Z.G. and P.S. wrote the manuscript. Z.G., T.C., N.Z., A.S., I.P., L.W., X.G., A.A., and R.L. contributed to the data acquisition and interpretation. P.S. supervised all work.

## Acknowledgements

The authors would like to thank Ms. Kathryn Nassar and Ms. Sandhya Bista for providing the *Borrelia burgdorferi* B31-A3 GFP strain and BSK-II medium, as well as technical support for *Borrelia burgdorferi* culture. The authors would like to thank Dr. Dinh-Tuan Phan for help with designing of algorithm for *Bb* quantification.

