## Supplementary information for "Dynamics of *Borrelia Burgdorferi* Invasion and Intravasation in a Tissue Engineered Dermal Microvessel Model"

**Supplemental Notes**

**Note S1**. Migration of *Bb* over long term of observation.

**Note S2**. Continuity of stationary, forward, and backward moving events in different environments.

**Note S3**. *Bb* migration in the perivascular region.

**Supplemental Figures**

**Figure S1**. *Borrelia Burgdorferi* local invasion model.

**Figure S2**. Distribution of forward, backward, and stationary moving events in different environments.

**Figure S3.** Comparison of *Bb* in the perivascular region with the cell body aligned parallel or perpendicular to the endothelium region of ECM.

**Figure S4**. Mechanisms of *Borrelia Burgdorferi* (*Bb*) transmigration.

**Supplemental movies**

**Movie S1**. *Bb* migration in the dermal microvessel model.

**Movie S2**. *Bb* migration in 1% methylcellulose solution.

**Movie S3**. Example of direct *Bb* transmigration.

**Movie S4**. Example of indirect *Bb* transmigration.

**Movie S5**. Example of *Bb* back and forth motion prior to transmigration.

**Movie S6**. Example of *Bb* back and forth motion at the ECM-microvessel interface prior to transmigration.

**Movie S7**. Example of transient tethering of the rear of the cell body of a *Bb* prior to intravasation.

**Supplemental Methods**

#### Cell Culture

*Fabrication of Tissue-Engineered Dermal Microvessels and Borrelia Inoculation*

#### Imaging

#### Bb tracking

#### Immune cell adhesion

#### Immunocytochemistry

#### Permeability

#### Statistics

### Supplementary Notes

#### **Note S1. Distribution of Bb following inoculation.**

*Bb* were injected into ECM approximately 900 µm from the midpoint of the dermal microvessel. The first *Bb* were observed beyond the microvessel within 1 h after inoculation. This is consistent with the predicted time (25 mins) to travel 900 µm, based on a net speed of 0.6 µm s^-1^. The distribution of *Bb* at 48 h post-inoculation was relatively symmetrical around the midpoint of the vessel, but with a small shift towards the downstream side (to the right in the images) (**Fig. 1g**), suggesting a small bias due to interstitial flow. In tracking individual *Bb* (average tracking time was 117 ± 138 s s) we observed no net bias perpendicular or parallel to the microvessel. However, a bias of 350 µm over 48 h corresponds to a net displacement of around 0.2 µm in 100 s (the average tracking time), which is below the imaging resolution (0.4 µm).

#### **Note S2. Comparison of forward, backward, and stationary events in different environments.**

In studies in 2D, which we recapitulated here, *Bb* exhibited three modes of motion: forwards, backwards, and stationary. In the bulk ECM and the perivascular region of dermal microvessels, we observed the same modes of motion. In all three environments (ECM, perivascular region, and 2D), the fraction of time that *Bb* were in the stationary was relatively small: 1.1% in ECM, 4.3% in the perivascular region, and 3.0% in 2D. The duration of stationary events was 1.00 ± 0.0 s in ECM, 1.05 ± 0.34 s in the perivascular region, and 1.07 ± 0.29 s in 2D (**Fig. S2h**).

The durations of forward and backward events were also measured (**Fig. S2d-g**). First, we consider forward and backward motion where forward motion is defined by θ ≤ 90˚ and backwards motion is defined by θ > 90˚. With this definition, *Bb* have shorter sustained forward motion both in bulk ECM (1.25 ± 0.54 s) and in the perivascular region (1.14 ± 0.38 s) compared to 2D (3.97 ± 6.20 s) (**Fig. S2h**). In particular, sustained forward motion was < 5 s in the ECM and perivascular region, while sustained forward motion of 5 - 25 s was relatively common in 2D (~20% of all forward motion), and occasionally was as long as 75 s (**Fig. S2d**). This trend was maintained when forward motion was defined by θ ≤ 30˚ (**Fig. S2e)**: the duration of forward motion was 1 - 3 s in ECM and the perivascular region, but typically 5 - 15 s, and sometimes as long as 25s, in 2D.

In contrast to forward motion, the difference between 3D and 2D was less striking for backwards motion. Persistent backward motion is defined by successive segments with θ > 90˚ and corresponds to back-and-forth motion. A significantly longer duration of backward motion was observed for *Bb* in ECM (2.23 ± 1.52 s) or in the perivascular region (2.24 ± 2.08 s) compared to in 2D (1.70 ± 1.21 s), however, the fold change was only about 1.3 (**Fig. S2h)**. Some relatively long backward moving events (10 - 20 s in duration) were observed for *Bb* in the perivascular region, but were not observed in 2D or ECM (**Fig. S2f**). When backwards motion was defined as θ >150˚, *Bb* in the perivascular region had significantly longer duration (2.03 ± 1.90 s) compared to 2D (1.26 ± 0.56 s, p < 0.0001) or ECM (1.73 ± 1.10 s, p < 0.01) although the fold change is only 1.17. These data indicate that θ > 150˚ accounts for a higher fraction of backward motion in the perivascular region compared to 2D and ECM, also providing an explanation for the lower net speed. The duration of backward moving events with θ > 150˚ (**Fig. S2g**) were typically 0 - 5 s long in 2D, while some of these events were as long as 5 - 10 s in the ECM and 10 - 20 s in the perivascular region.

To measure how often *Bb* changed their migration direction, we also analyzed the frequency of direction reversal events (**Fig. S2i)**. Independent of the definition of a reversal event (i.e. θ > 90˚, 120˚, 150˚, or 170˚), *Bb* exhibited more reversals in 3D (ECM or perivascular region) compared to 2D (methylcellulose solution). In addition, *Bb* had a higher frequency of reversals with θ > 150˚ in the perivascular region compared to the ECM (**Fig. S2i**), which is consistent with the observation of increased back and forth motion in the perivascular region than in ECM. (**Fig. S2e,f**). In the literature, a reversal rate of 18.89 ± 5.08 min^-1^ has been reported for *Bb* in 1% methylcellulose solution (1). While the definition of a reversal event was not reported, the value is similar to our results for θ > 90˚ (**Fig. S2i)**.

Together, these results support the following conclusions. (1) The sustained duration of forward motion in 2D compared to 3D explains the higher net displacement speed (compared to 3D). The lower instantaneous speed in 2D is due to the viscosity of the methylcellulose solution. (2) The higher fraction of stationary events and the increased number of reversals in the perivascular region compared to bulk ECM explains the lower instantaneous and net displacement speeds (compared to bulk ECM).

#### **Note S3. Bb migration in the perivascular region**

In the perivascular region, the endothelium provides an obstacle to motion. To assess whether there was any influence of *Bb* orientation on migration in the perivascular region, we analyzed 2,051 segments from 5 *Bb*. The *Bb* in these segments were classified as either parallel to the endothelium, perpendicular to the endothelium, or neither. Parallel refers to the case where the whole of the *Bb* cell body is aligned parallel to endothelium (n = 1,023 segments). Perpendicular refers to the case where one end of the *Bb* cell body is close to or touching the endothelium and the cell body is at a distinct angle to the endothelium (denoted for convenience as perpendicular) (n = 313 segments). Intermediated cases where part of the cell body was parallel to the endothelium and part was at a distinct angle were excluded from the analysis. In addition, *Bb* undergoing transendothelial migration were excluded from analysis. We compared the migration behavior of *Bb* in the perivascular region in these two in states (**Fig. S3**). *Bb* parallel to the endothelium had lower instantaneous speed (2.62 ± 1.86 µm s^-1^) compared to perpendicular to the endothelium (4.37 ± 2.89 µm s^-1^) (**Fig. S3a**). The fraction of stationary events was higher for *Bb* parallel to the endothelium (8.5%) compared to perpendicular (1.9%) (**Fig. S3b**). The only stationary events > 1 s (t = 2 s and 4 s) occurred for *Bb* parallel to the endothelium, while all stationary events for *Bb* perpendicular to the endothelium was ≤ 1 s (**Fig. S3c**). These results suggest that the decrease of instantaneous speed and increased fraction of stationary events for *Bb* parallel to the endothelium could be associated with interactions with the basement membrane or endothelium, although there was no significant increase in the duration of stationary events.

### Supplemental Figures

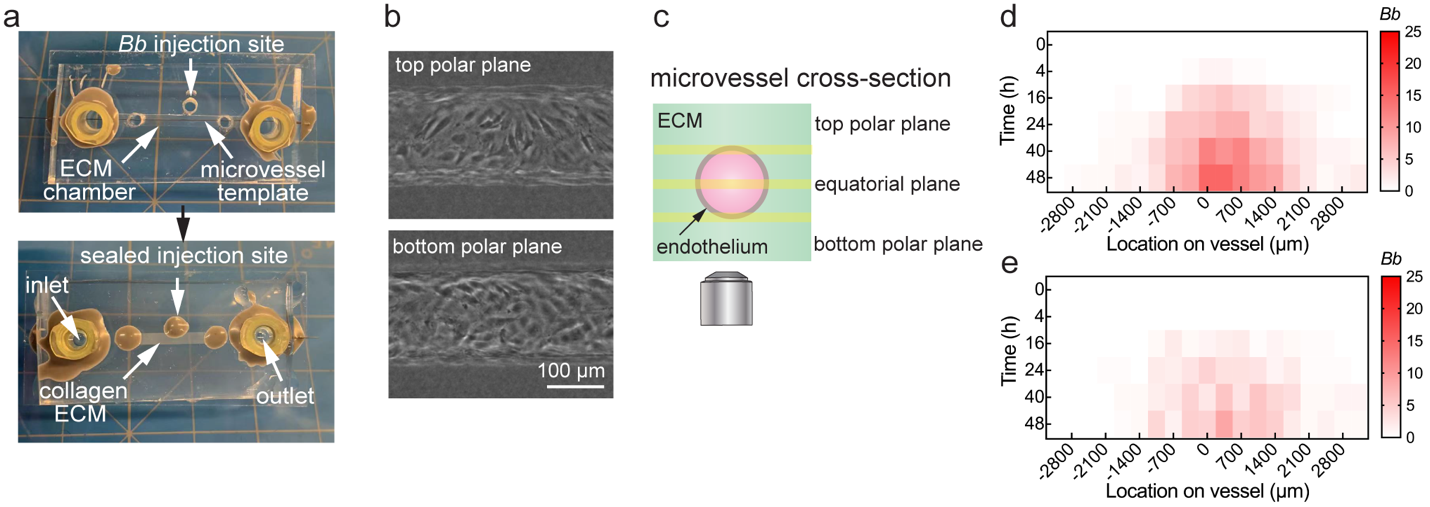

**Figure S1***.* *Borrelia Burgdorferi* local invasion model.

(a) Photograph of the dermal microvessel platform. (Left) Platform with template rod prior to injection of ECM. (Right) After injection of collagen I matrix and removal of template rod. The injection port is sealed with elastomer after injection of *Bb* to prevent fluid leakage due to the transmural pressure.

(b) Phase contrast images of the top and bottom polar planes of a microvessel showing a confluent monolayer of dermal microvascular endothelial cells 48 h after seeding.

(c) Schematic illustration showing imaging locations.

(d) Distribution of *Bb* in the perivascular region anterior to the inoculation site.

(e) Distribution of *Bb* in the perivascular region posterior to the inoculation site.

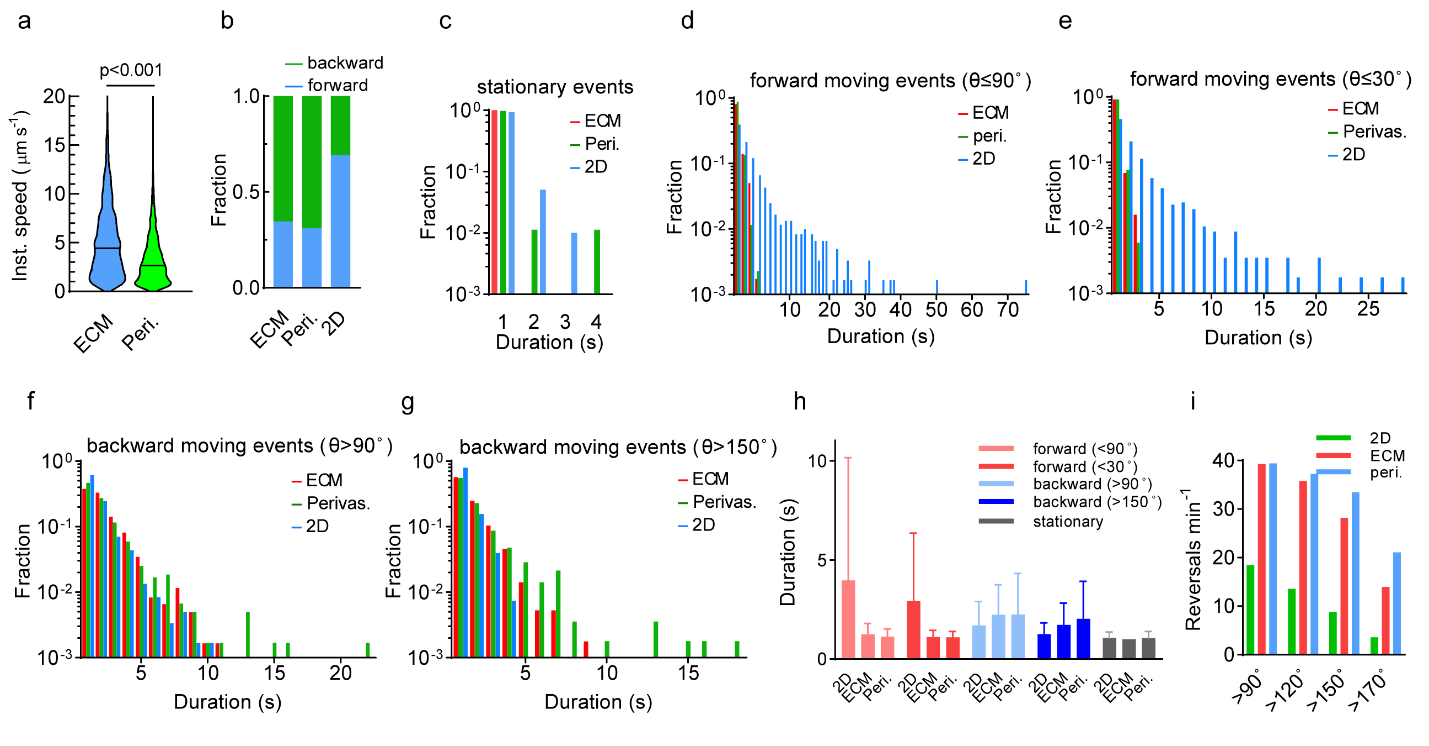

**Figure S2***.* Distribution of forward, backward, and stationary moving events in different environments.

(a) The instantaneous speed of *Bb* in the ECM and in the perivascular region. ECM: 2,104 segments, 20 *Bb*. Perivascular region: 2,035 segments, 5 *Bb*.

(b) Fraction of forward and backward events in the ECM, the perivascular region, or in 2D. Forward events: θ ≤ 90˚. Backwards events: θ > 90˚.

(c) Distribution of the duration of stationary events in the ECM, the perivascular region, or in 2D. ECM: 23 segments, 20 *Bb*. Perivascular region: 87 segments, 5 *Bb*. 2D: 106 segments, 18 *Bb*.

(d) Distribution of duration of forward moving events with θ ≤ 90˚ in the ECM, the perivascular region, or in 2D.

(e) Distribution of duration of forward moving events with θ ≤ 30˚ in the ECM, the perivascular region, or in 2D.

(f) Distribution of duration of backward moving events with θ > 90˚ in the ECM, the perivascular region, or in 2D.

(g) Distribution of duration of backward moving events with θ > 150˚ in the ECM, the perivascular region, or in 2D.

(h) Duration of forward moving events (θ ≤ 90˚ or 30˚), backward moving events (θ > 90˚ or 150˚), and stationary events in 2D, the ECM, and in the perivascular region. Bars: mean±SD.

(i) Number of reversals per minutes for *Bb* in 2D, the ECM, or the perivascular region with backwards events defined by (θ > 90˚, 120˚, 150˚, or 170˚)

(b and d-i) ECM: 2,081 segments, 20 *Bb*. Perivascular region: 1,948 segments, 5 *Bb*. 2D: 3,470 segments, 18 *Bb*.

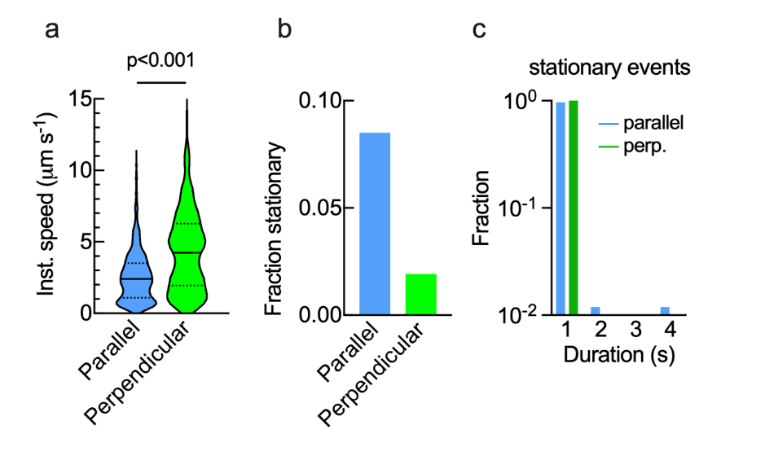

**Figure S3.** Comparison of *Bb* in the perivascular region with the cell body aligned parallel or perpendicular to the endothelium. (a) Instantaneous speed of *Bb* in the perivascular region. (b) Fraction of stationary events (of all events) in the perivascular region. (c) Fraction of the duration of stationary events in the perivascular region.

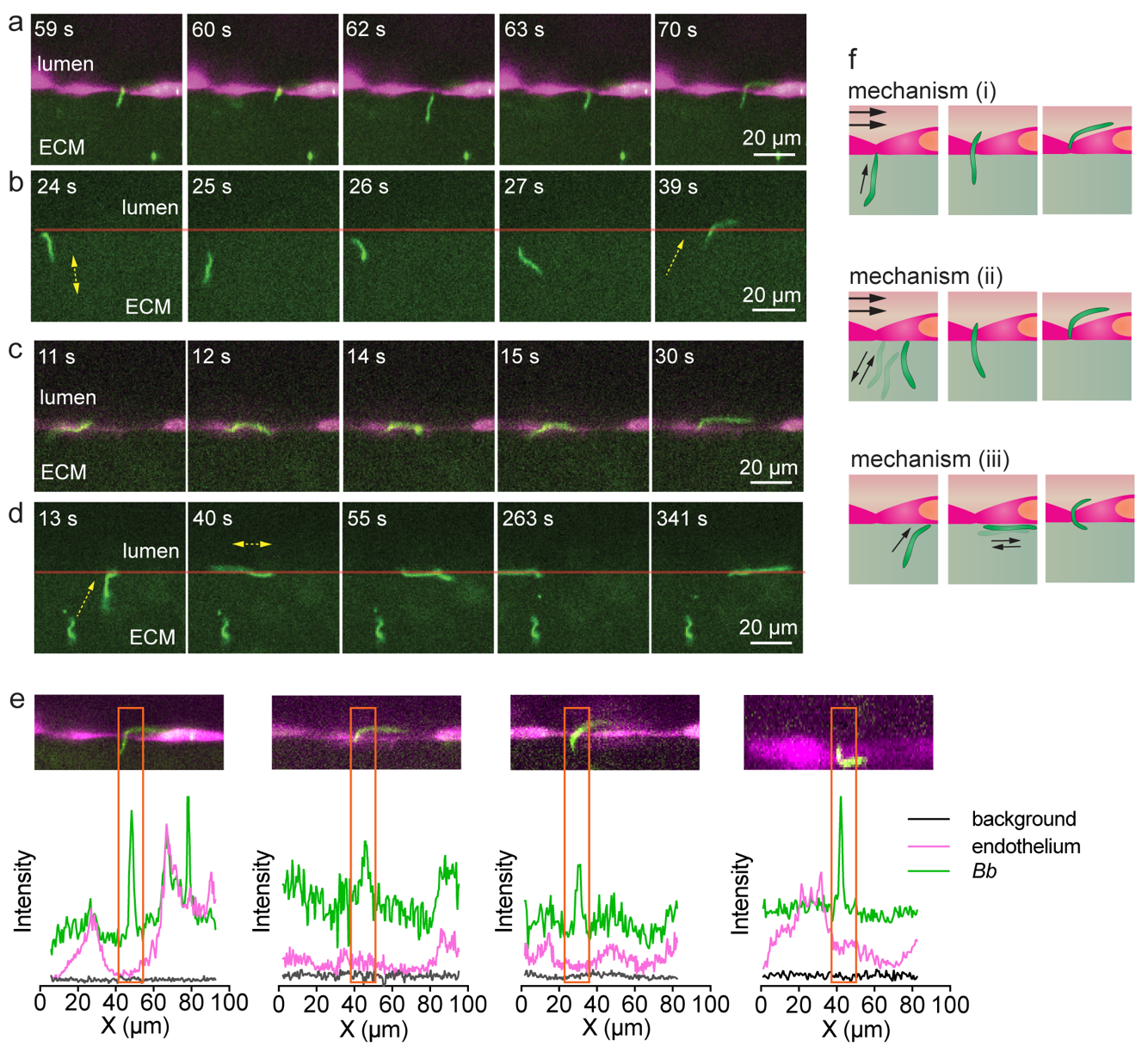

**Figure S4**. Mechanisms of *Borrelia Burgdorferi* (*Bb*) transmigration.

(a) Example of indirect intravasation, where a *Bb* arrives at the microvessel and then undergoes back and forth motion in the ECM perpendicular to the microvessel prior to transmigration at a cell-cell junction.

(b) Example of indirect intravasation, where a *Bb* arrives at the microvessel and then undergoes back and forth motion in the ECM perpendicular to the microvessel prior to transmigration. In this example, the endothelial cells are not labeled. The red line indicates the position of the endothelium (from phase images).

(c) Example of indirect intravasation, where a *Bb* arrives at the microvessel and then undergoes back and forth motion in the ECM parallel to the microvessel prior to transmigration at a cell-cell junction.

(d) Example of indirect intravasation, where a *Bb* arrives at the microvessel and then undergoes back and forth motion in the ECM parallel to the microvessel prior to transmigration. The endothelial cells are not labeled. The red line indicates the position of the endothelium (from phase images).

(e) Fluorescence intensity profiles of *Bb* (green) and endothelial cells (magenta) along the endothelium at a location where a *Bb* undergoes transmigration. The peak of the *Bb* fluorescence (green) coincides with the minimum fluorescence (magenta) of the endothelial cells (i.e. the location of the cell-cell junctions).

(f) Schematic illustration of the mechanism of direct and indirect intravasation through a cell-cell junction. (i) A *Bb* arrives at the endothelium at a cell-cell junction and immediately undergoes transmigration. (ii) A *Bb* arrives at the endothelium, contacts a cell body, and then undergoes back and forth motion perpendicular to the microvessel prior to locating a cell-cell junction and undergoing transmigration. (iii) A *Bb* contacts an endothelial cell body and undergoes back and forth motion parallel to the interface prior to locating a cell-cell junction and undergoing transmigration.

### Supplemental movies

#### Movie S1

*Bb* migrate in the collagen I ECM of the microvessel model over 15 min. *Bb* stays in focus for ~100s and z-axis migration will result in it disappear in the field of view. Analysis of migration path, speed, angle of vectors, intravasation in the microvessel model are obtained from movies with scale of field of view and image duration similar to movie S1. Scale bar: 100 μm.

#### Movie S2

Bb migrate in the 1% methylcellulose solution over 15 min. Analysis of migration path, speed, angle of vectors in the solution are obtained from movies with scale of field of view and image duration similar to movie S2. Scale bar: 100 μm.

#### Movie S3

*Bb* (indicated by white arrow) transmigrated immediately after contacting endothelial cell-cell junction. Scale bar: 25 μm.

#### Movie S4

*Bb* (indicated by white arrow) touched the endothelial cell body, then migrated away to the cell-cell junction region and transmigrated; another *Bb* (indicated by orange arrow) touched the endothelial cell body for several times and failed to intravasate, and disappeared in the field of view eventually. Scale bar: 25 μm.

#### Movie S5

When touching endothelium, *Bb* (indicated by white arrow) moved back and forth in ECM for a few rounds before transmigration. Scale bar: 25 μm.

#### Movie S6

When touching endothelium, *Bb* (indicated by white arrow) moves back and forth along the vessel-ECM interface for a few rounds before transmigration. Scale bar: 25 μm.

#### Movie S7

*Bb* moved back and forth on ECM-endothelium interface (indicated by white arrow), and at the time point that the orange arrow appeared, it became stationary with only one end tethering to the endothelium, indicating the entire *Bb* transmigrated and exposed to flow. Scale bar: 25 μm.

### Supplemental Methods

#### Cell Culture

Adult human dermal microvascular endothelial cells (HDMECs) (Lonza, CC-2543) were used to generate tissue-engineered dermal microvessels. HDMECs were cultured in EGM™-2 Endothelial Cell Growth Medium-2 (EC media; Lonza) from passage 3 to 5, using 0.25% Trypsin-EDTA (ThermoFisher). Note that GA-1000 provided in the supplementary kit (EGMTM-2 MV microvascular endothelial cell growth medium SingleQuots^TM^ supplements (CC-4147)) was replaced with gentamicin at a final concentration of 30 μg mL^-1^, as the *Bb* strain used in the study is gentamicin resistant. The *Bb*-GFP (B31-A3 GFP strain) was cultured in BSK medium at 34 ˚C with 5% CO_2_. The *Bb-GFP* used throughout the study were at passage 4 to avoid loss of endogenous plasmids caused during passing.

*Fabrication of Tissue-Engineered Dermal Microvessels and Borrelia Inoculation*

The platform for the dermal microvessel was based on our previous work(2, 3), with the addition of a side port for *Bb* inoculation (**Fig. 1a**). 7 mg mL^-1^ neutralized collagen I solution (Corning™ collagen I, high concentration, rat tail) was carefully injected into the chamber of the device housing (**Fig. S1b**). in the room temperature and then gelled for 20 - 30 min at 37 ˚C. Next, the template rod (**Fig. S1a**) was removed to create a 150 μm channel. The side opening was filled with PBS to keep collagen hydrated and then sealed with silicone elastomer (Slygard^TM^ 164, Dow Corning). The collagen gel was then crosslinked by perfusion with 20 mM genipin (Wako Chemicals USA), for 2 hours, and then perfused with PBS (cat. no. 10010072, Gibco™), and perfused with 2~4 μL high concentration dermal microvascular endothelial cell (HDMEC). Cell seeding was performed by introducing HDMECs into the channel for 10 min under static conditions at a cell density of ~ 4 × 10^7^ mL^-1^ in EC media following previously reported protocols (2). Cell adhesion and spreading was allowed to proceed under static conditions for 30 min. Following formation of a confluent monolayer (~ 12 h), the microvessels were perfused with endothelial cell medium (EGM-2) under shear stress of ~4 dyne cm^-2^ for 2 days to allow maturation. Phase contrast images (**Fig. 1b and Fig. S1c-e**) show the formation of a confluent monolayer.

After maturation, *Bb* were inoculated in the side port of the dermal microvessel. A syringe with ~5 × 10^8^ *Bb* in 20 μL BSK-2 medium with a 23G needle was inserted into the elastomer covering the side port and the *Bb* gently injected. The injection site was then quickly sealed with freshly prepared silicone elastomer to minimize flow from the microvessel to the side port due to the transmural pressure.

For 2D “swimming” experiments, ~1 × 10^7^ *Bb* in 40 - 50 μL 1% methylcellulose dissolved in medium (50% BSK medium + 50% PBS) was injected into the empty chamber of a microfluidic device (**Fig. S1**) and maintained in an incubator for at least 2 hours before imaging. Imaging was performed using a confocal microscope at 20× magnification (for details see *Imaging* section).

#### Imaging

Confocal images were acquired at 20× magnification on a swept field confocal microscope system (Prairie Technologies) and illumination was provided by an MLC 400 monolithic laser combiner (Keysight Technologies). Images of the whole microvessel were assembled from 100 images of approximately 7 mm segments along the length. Images were recorded at 0 h, 4 h, 24 h, 40 h, and 48 h after inoculation. These images were used to determine the number of *Bb* in the vicinity of microvessel (˂ 20 μm), which were counted manually.

Time lapse fluorescence images were acquired at 20× magnification on a swept field confocal microscope system (Prairie Technologies) for 1 h with image rate of 1s/frame, both in the microvessel model and in methylcellulose solution. Imaging was performed in an environmental chamber at 37 ˚C with > 90 % RH, and 5 % CO_2_. Movies were obtained at 24 h or 48 h after *Bb* inoculation. Before and after fluorescence imaging, phase contrast images were recorded to ensure microvessel integrity. To visualize the endothelium and *Bb*, in some experiments HDMECs were stained with CellTracker Deep Red (Thermo Fisher Scientific, C34565) in a T-25 flask according to product protocols 1 day prior to seeding in the device.

#### Bb tracking

Tracking of individual *Bb* was performed in Image J. The center of mass of a *Bb* cell body visible in an image was determined manually and used to represent the position of *Bb*. Only *Bb* that remained within the focal plane for more than 20 seconds were included in tracking statistics. Specifically, *Bb* were included if the fluorescence intensity was substantially higher than the surrounding ECM (average *Bb* intensity more than 1.5 times higher than the average intensity from 5 randomly selected regions with no *Bb*), and a significant displacement between frames was observed. The coordinates of the center of mass at time point (t) were defined as (x_t_, y_t_), and hence the length of the displacement vector (d_t_) is given by:

$d_{t}=\sqrt{\left( y_{t}-y_{t-1} \right)^{2}+\left( x_{t}-x_{t-1} \right)^{2}}$

The instantaneous speed is d_t_/Δt where the time between images Δt = 1s. Therefore, the instantaneous speed of the tracked *Bb* is:

$s_{t}=\frac{\sqrt{\left( y_{t}-y_{t-1} \right)^{2}+\left( x_{t}-x_{t-1} \right)^{2}}}{\Delta t}$

The net speed of the tracked *Bb* is determined from the net displacement between the first and last frames and the tracking time:

$$s\left( net \right)=\frac{\sqrt{\left( y_{t}-y_{0} \right)^{2}+\left( x_{t}-x_{0} \right)^{2}}}{n\Delta t}$$

where n is the number of segments along a migration path and the total time of the migration path is n∆t.

The angle θ (degrees) between two consecutive displacement vectors is given by:

$$\theta=\frac{180}{\pi}acos\frac{(y_{t}-y_{t-1})(x_{t}-x_{t-1})+(y_{t-1}-y_{t-2})(y_{t-1}-y_{t-2})}{\sqrt{\left( y_{t}-y_{t-1} \right)^{2}+\left( x_{t}-x_{t-1} \right)^{2}} \sqrt{\left( y_{t-1}-y_{t-2} \right)^{2}+\left( x_{t-1}-x_{t-2} \right)^{2}}}$$

In most cases, *Bb* motion was defined as forward when θ ≤ 90˚, backward when θ > 90˚, and stationary when d ≤ 0.4 μm (~2% of the typical length of a single *Bb* cell body). When *Bb* were stationary those frame(s) were excluded from the calculation of θ. The duration of the different modes of motion (forward, backward, and stationary) were defined as the number of successive vector segments along the migration path that satisfied with the criteria outlined above. The resolution of analysis was defined by the image collection rate (1 s).

The distance of *Bb* to the microvessel were calculated from each frame using ImageJ. The microvessel boundary was determined from a linear least squares fit (y=ax+b) to 5 points along boundary. At any time point (t) the distance between the *Bb* (x_t_,y_t_) and the microvessel was calculated from |y_t_ – (ax_t_+b)|.

For analysis we define the perivascular region as the region within 20 µm of the endothelium, i.e. approximately within the average length of the *Bb* cell body. The region > 20µm from the microvessel is denoted as bulk ECM. The tracking of *Bb* during intravasation was performed using the same methods as described above, during imaging at the equatorial plane of the microvessel.

#### Immune cell adhesion

To assess immune cell adhesion, microvessels were perfused with HL-60 (neutrophil-like) and THP-1 (monocyte-like) cells. At 48 hours after inoculation with 1×10^7^ *Bb* or vehicle (BSK medium), microvessels were perfused with immune cells for 10 min. Human HL-60 (ATCC, CCL-240) and THP-1 (ATCC, TIB-202) cells were cultured in RPMI-1640 medium (Thermo Fisher Scientific, 11875093) supplemented with 10% fetal bovine serum (Sigma, F4135) and 1% penicillin-streptomycin (Thermo Fisher Scientific, 15140122). Both HL-60 and THP-1s were maintained in liquid nitrogen and thawed 24 hours prior to use. Before each experiment, HL-60 or THP-1s were labeled with dyes according to product protocols for live cell tracking. HL-60 were labelled with Calcein-AM (Thermo Fisher Scientific, C1430). THP-1 were labelled with CellTracker Deep Red (Thermo Fisher Scientific, C34565). After washing the cells twice with PBS, both HL-60 and THP-1s were resuspended at a final concentration of 1 × 10^6^ cells mL^-1^ in EGM-2 medium and 70 µL of this suspension (approximately 70,000 HL-60 and 70,000 THP-1) was perfused through the device for 10 minutes under low shear stress (~0.2 dyne cm^-2^). Afterwards, non-adherent cells were removed by perfusing microvessels with EGM-2 medium to for 20 minutes. Adherent HL-60 and THP-1s were manually counted in each device, and normalized to the area of microvessel.

#### Immunocytochemistry

Immunocytochemistry was performed on microvessels 48 h after *Bb* inoculation. Microvessels were perfused and washed with DPBS (Invitrogen, 14080055) for 10 minutes, and then fixed with 3.7% paraformaldehyde solution overnight at 4 ˚C. Microvessels were then stained by: (1) blocking and permeabilizing with blocking solution composed of 0.2% Triton-X 100 (Sigma-Aldrich, T9284) and 10% normal goat serum (Cell Signaling Technology, 5425S) overnight at 4 ˚C, (2) perfusing with primary antibody solution for 6 h at room temperature, (3) washing with blocking solution overnight at 4 ˚C, (4) perfusing with secondary antibody solution for 30 min at room temperature, and (5) washing with blocking solution overnight at 4 ˚C before imaging. ICAM-1 was stained with anti-human ICAM-1 (Abcam, ab2213), and F-actin was stained with Alexa Fluor™ Plus 647 Phalloidin (Thermo Fisher, A30107) without blocking buffer perfusion. Confocal z-stacks (0.4 µm in thickness) were acquired at 40× magnification on a swept field confocal microscope system (Prairie Technologies) and illumination was provided by an MLC 400 monolithic laser combiner (Keysight Technologies). Reconstructions of microvessels were assembled from approximately 400 slices. To quantify the expression level of ICAM-1 (**Fig. 4g**), maximal intensity projection fluorescence images of z stacks of 400 slices were analyzed in Image J after background subtraction. Reported data were normalized to the average fluorescence intensity of microvessels (n = 7) from the control group (vehicle).

#### Permeability

To assess changes in permeability, at 48 hours after inoculation, microvessels with 1×10^7^ *Bb* or vehicle (BSK medium) were perfused with 2 µM Alexa Fluor647-conjugated 2 MDa dextran (Thermo Fisher Scientific, D22914) in EGM-2. Microvessels were then imaged (Nikon Eclipse TiE) at 10× magnification and maintained in a live cell chamber at 37 ˚C. Epifluorescence illumination was controlled by X-Cite 120LEDBoost (Excelitas Technologies). Phase contrast images (8107 μm × 664 μm) were acquired every 30s at both planar and equatorial planes of the microvessel; fluorescence images were acquired every 30s at the microvessel equatorial plane. Microvessels were imaged for 2 minutes before and 5 minutes following perfusion with the fluorescent solutes.

To calculate permeability, images were cropped (ImageJ, NIH) and sectioned into 10 regions of interest, each 0.81 mm × 0.66 mm. The integrated pixel density was plotted over time for each ROI and the permeability (cm s^-1^) was calculated using P=(r/2)(1/∆I)(dI/dt), where r is the vessel radius, ∆I is the increase in fluorescence intensity upon initiation of perfusion of the solute, and dI/dt is the rate of fluorescence intensity increase as the solute permeates into the collagen gel. The average permeability was calculated via linear least squares fits of dI/dt over 6 frames (10 minutes).

#### Statistics

Statistical analyses were conducted using Prism (GraphPad ver. 9). All experimental values are reported as mean ± standard deviation (S.D.). A student’s unpaired t-test (two-tailed with unequal variance) was used for comparison of two groups. Differences were considered statistically significant for p < 0.05, with the following thresholds: * p < 0.05, ** p < 0.01, *** p < 0.001. Linear regression was conducted using least squares fitting with no constraints, and an F-test was used to determine if linear regression produced a statistically significant non-zero slope.
